# AdjuSST: An Adjustable Surface Stiffness Treadmill

**DOI:** 10.1101/2024.03.25.586685

**Authors:** Mark Price, Dominic Locurto, Banu Abdikadirova, Meghan E. Huber, Wouter Hoogkamer

## Abstract

Humans have the remarkable ability to manage foot-ground interaction seamlessly across terrain changes despite the high dynamic complexity of the task. Understanding how adaptation in the neuromotor system enables this level of robustness in the face of changing interaction dynamics is critical for developing more effective gait retraining interventions. We developed an adjustable surface stiffness treadmill (AdjuSST) to trigger these adaptation mechanisms and enable studies to better understand human adaptation to changing foot-ground dynamics. The AdjuSST system makes use of fundamental beam-bending principles; it controls surface stiffness by controlling the effective length of a cantilever beam. The beam acts as a spring suspension for the transverse endpoint load applied through the treadmill. The system is capable of enforcing a stiffness range of 15-300kN/m within 340 ms, deflecting linearly downwards up to 10 cm, and comfortably accommodating two full steps of travel along the belt. AdjuSST offers significant enhancements in effective walking surface length compared to similar systems, while also maintaining a useful stiffness range and responsive spring suspension. These improvements enhance our ability to study locomotor control and adaptation to changes in surface stiffness, as well as provide new avenues for gait rehabilitation.

## I. INTRODUCTION

During walking or running, the human nervous system must control the many degrees of freedom in the musculoskeletal system while also managing the dynamic and discontinuous interaction between the feet and the ground. Nevertheless, people can seamlessly manage this task in their daily lives, across often abruptly varying terrains. Understanding the mechanisms that humans exploit to manage these changes will provide promising directions to help improve gait in neurologically impaired individuals. When we can access the neural mechanisms for changing motor control strategy, we should be able to shape desired changes in neurologically impaired motor control, such as stroke-induced hemiparesis. However, it remains an challenge to identify these mechanisms for change.

It is possible to observe the ways in which the nervous system adapts to locomotion challenges by suddenly changing the nature of the human-environment interaction while recording the behavioral response [1]–[3]. Observing the nature of this response can expose underlying patterns that humans rely on for gait and balance, not only for robustness to sudden change, but also for effective management of the new condition [4]–[7]. One such method that has been used for studying and exploiting neuromotor adaptation for rehabilitation is the use of split belt treadmills [8], [9]. In split belt treadmill studies, belt speeds are altered asymmetrically, resulting in spatiotemporal adaptations over time [10]. While split belt treadmills have demonstrated the ability to affect step length asymmetry on long time scales after training for people after stroke [11], kinetic measures such as weight-bearing and propulsion asymmetry remain resistant to long-term correction [12]–[15] and to transfer to overground gait [11], [14]. This gap is critically important for rehabilitation outcomes, because asymmetric weight-bearing is correlated with increased risk for injury [16] and developing additional impairments, such as knee osteoarthritis [17]–[19].

We posit that split belt treadmill training has been more successful in correcting kinematic over kinetic measures because it imposes kinematic constraints as a means of perturbing gait. A device that alters the interface dynamics between the foot and the ground may be more effective in eliciting motor adaptations in gait kinetics. This paper presents the design of an adjustable surface stiffness treadmill (AdjuSST) and validation of its capability to create our proposed conditions for future human experiments. The primary contribution of this work is its novel design relative to existing adjustablt stiffness treadmills enabling it to meet performance goals specific to our intended use case: within-step stiffness change under load, sufficient effective length to allow minor walking speed fluctuations and to avoid restricting natural gait mechanics or responses to perturbations, and consistent vertical stiffness along that length.

This paper is organized as follows: 1) we begin by describing the prior art in the controllable surface stiffness treadmill design space and defining the solution required for our application, then we 2) present the full mechatronic design of the AdjuSST system, 3) present our methods and findings for characterizing the performance of the system, and finally 4) discuss our findings, compare them with the prior work, and provide insights for future design of similar systems.

## II. PRIOR ART

### A. Adjustable stiffness treadmills

We are aware of two treadmills with adjustable surface stiffness that have been created by other researchers [20], [21]. The first published design (Variable Stiffness Treadmill, or VST) uses a moving fulcrum to adjust the mechanical advantage of a lever arm attached to a set of coil springs and rotates about an axis located at the front of the treadmill [20]. The VST allows for a high range of stiffnesses and is capable of making maximal stiffness changes in under 250 ms, allowing it to change stiffness within one step with a comfortable margin. The design offsets the weight of the treadmill belt and platform with a counterweight suspended in front of the rotation axis, ensuring that the belt always returns to the horizontal position regardless of the stiffness setting.

The VST has been successfully implemented for human participants research. Among other findings, single-step stiffness perturbations have been shown to increase plantarflexor activity in participants affected by stroke-induced hemiparesis [22], and longer exposures to asymmetric stiffness appear to result in changes to step length and muscle activity after the exposure in healthy participants [23].

However, the design has two characteristics that do not support our desired experiments. First, by controlling torsional stiffness about an axis at the front of the treadmill, the apparent vertical stiffness of the ground depends on the distance of the foot from the rotation axis. As the center of pressure of the ground reaction force translates along the belt, the moment arm about the rotation axis increases. Therefore, either constant active control of the fulcrum position in response to sensing of the foot position is required, or the vertical stiffness decreases with distance from the front of the treadmill, changing the nature of the perturbation.

The second characteristic is caused by the first: the allowable step length and foot position on the belt is limited, with little margin for the walker to fluctuate their walking speed or foot placement strategy. By employing a torsional stiffness for the belt, the system is placed under significantly more load if the walker is allowed to drift toward the back of a longer platform. While the effective walking surface length is not reported, the VST uses an 80 cm long force sensing mat beneath the belt to detect ground reaction forces, which is near the upper range of a typical walking step length, depending on the height and speed of the walker. The belts accommodate slightly more than one step length, which can be observed from figures which include experimental participants [20], [23].

An adjustable stiffness treadmill design with linear vertical deflection (Treadmill with Adjustable Surface Stiffness, or TwAS) has also been presented [21]. It uses the same principle of a controllable fulcrum point, but transfers the load through a scissor linkage to the treadmill, which functions as a linear motion constraint. While this addresses the first limitation, any increase in the length of the walking surface from the VST is not reported and is not apparent from photographs. Furthermore, while a similar mechanism to the VST is used to adjust the stiffness, the maximum achievable stiffness is significantly reduced, from 2000 kN/m for the VST to 40 kN/m for the TwAS. For comparison, this is lower than the minimum stiffness tested in experiments investigating the effect of surface stiffness on running energetics [24]. While the linkage mechanism appears to be effective in constraining the treadmill to deflect vertically, the structural rigidity is low enough that it may interfere with attempts to measure the effect of changing surface stiffness relative to a rigid treadmill.

### B. Alternative mechanisms

A potential solution to creating an adjustable stiffness mechanism with high structural rigidity may be found in adjustable-length leaf spring designs [25]–[27]. Operating on a similar principle to the adjustable lever fulcrum employed by the VST and TwAS, variable length leaf springs have the advantage of being more compact as the structural function of the lever is performed by the spring itself, thereby allowing a short and direct path from the load to the ground when the length of the leaf spring is reduced to near zero. Variable length leaf spring designs have been successfully implemented for walking loads in variable stiffness prosthetic feet [28], [29]. Additionally, the force required to maintain a set stiffness with a variable-length leaf spring has been demonstrated to be bounded by the spring parameters to be low relative to the load applied to the spring [25], [30], allowing the actuation requirements to be confidently predicted and for control of the stiffness change to be highly precise and resistant to disturbance loads applied to the spring.

Therefore, we innovated upon prior controllable stiffness treadmill designs to use a variable-length leaf spring mechanism. In the next section, we present the full system design, with additional innovations to make the following changes from existing designs: (1) the adjustable stiffness must be linear stiffness constrained to the vertical axis, (2) the structure must have rigidity sufficient to present a high-stiffness surface of at least *>*100 kN/m, and (3) the effective walking surface length must be at least 2 step lengths to allow the walker freedom to adjust their gait strategy in response to the perturbation.

## III. DESIGN

### A. Overview

The design of the full variable stiffness treadmill system is illustrated in Fig. 1. The system comprises five subassemblies: a narrow, low-inertia treadmill; a leaf spring variable stiffness suspension; a preload linkage mechanism; a structural frame with vertical linear constraints; and an offboard motor assembly coupled to the treadmill via universal joints. The treadmill is designed to translate vertically up and down along its vertical rails, supported by the bending stiffness of a cantilevered sheet of spring steel, the effective length of which is modified via a servo-controlled rack and pinion. The treadmill geometry, offboard motor, and spring-based weight compensation (as opposed to an inertial counterweight) contribute toward minimizing the inertia of the treadmill, increasing the transparency of the device and increasing the usable range of walking speeds. This system is located inside a motion capture volume with 12 cameras (Qualisys, Gothenburg, Sweden) and mounted directly adjacent and parallel to a separate, rigidly mounted treadmill [31]. The system is designed to function as a dual belt treadmill, with each belt controlled independently with a separate motor, to allow the future study of behavioral response to imposed ground stiffness asymmetry.

**Fig. 1.**
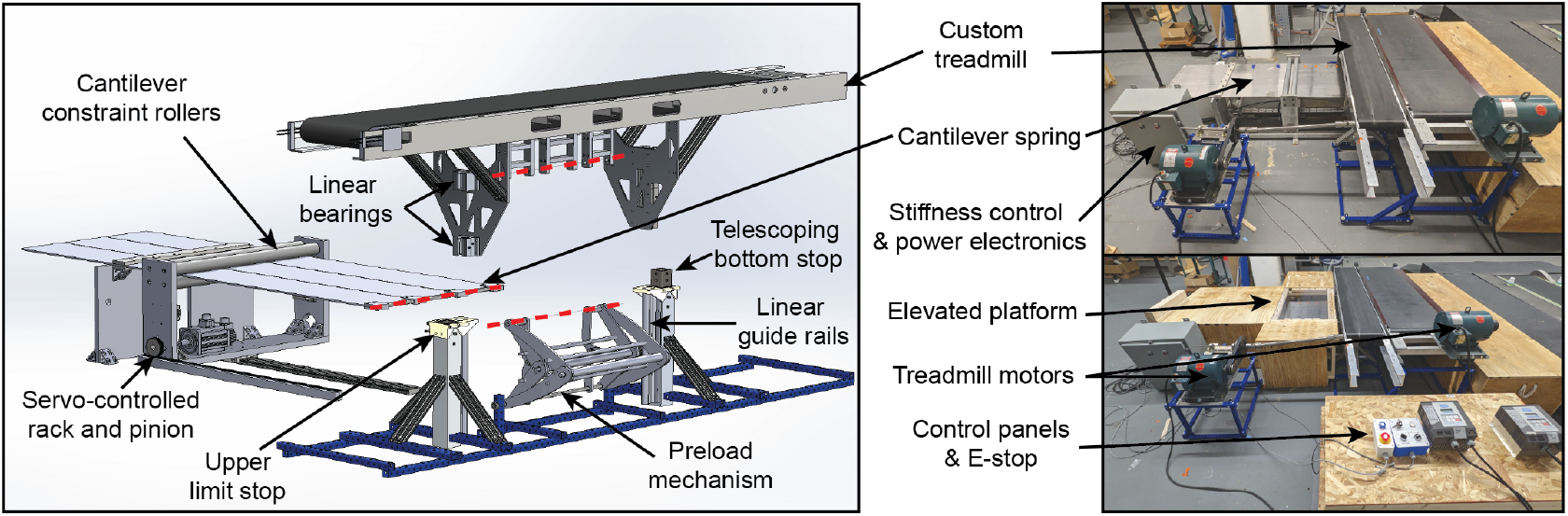
The AdjuSST system with labeled components. The dashed line in the exploded assembly indicates the coupling axis for the treadmill, adjustable stiffness mechanism, and preload mechanism. Note that the elevated platform has a window to allow the movement of the rollers to be observed. This window is covered with impact-resistant transparent plastic and is safe to walk on.

### B. Adjustable Stiffness Mechanism

The key principle of the adjustable stiffness mechanism is illustrated in Fig. 2A. Vertical stiffness (*dF/dd*) is a function of the effective cantilever length *L*_*e*_ of the leaf spring

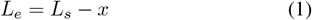

where *L*_*s*_ is the unmodified length of the beam and *x* is the distance of the cantilever constraint from the base of the beam. Variable stiffness mechanisms with this operating principle have been developed and accurately characterized for rotary joints [25]–[27]. This section extends the prior work to characterize the length-dependent stiffness behavior of a vertically displacing load with a separate torsion spring-based preload mechanism.

**Fig. 2.**
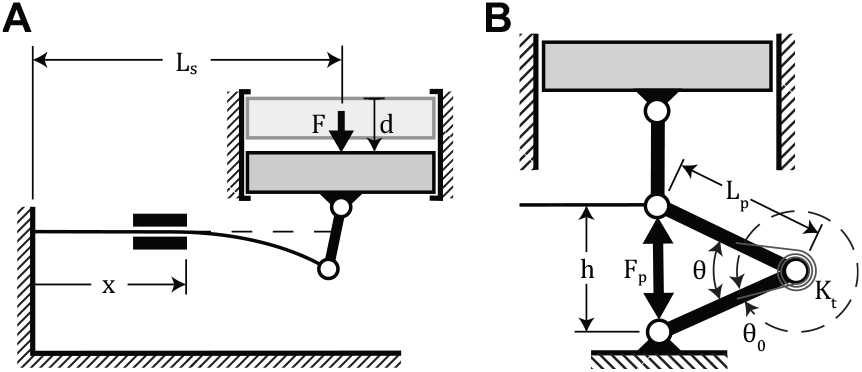
**A:** Parameters and geometry defining the adjustable stiffness spring mechanism. Note that deflection proportions are exaggerated for increased readability. **B:** Parameters and geometry for the unloaded deflection compensation mechanism.

Assuming a small deflection angle, the force-deflection relationship of the leaf spring can be modeled as a cantilever beam:

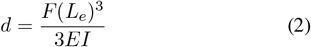

where *d* and *F* are the deflection of and force applied to the beam tip normal to its longitudinal axis, respectively, *E* is the Young’s modulus, and *I* is the area moment of inertia of the beam. Rearranging Eqn. 2 to solve for *F* and differentiating with respect to the deflection *d*, the stiffness of the beam

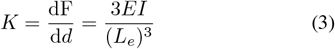

is proportional to the inverse cube of the effective beam length *L*_*e*_.

### C. Preload Mechanism

Because our mechanism is loaded by the treadmill and the unsupported weight of the spring, we add a distributed load term to Eqn. 2 and a point load to the beam tip to account for the added downward force:

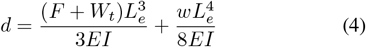

where *w* is the load per unit length associated with the weight of the exposed length of the spring, and *W*_*t*_ is the weight of the treadmill. After rearranging Eqn. 4 to solve for *F*

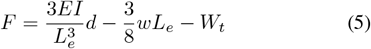

it becomes apparent that stiffness *K* remains defined by Eqn. 3 because the added weight terms are not dependent on *d*. However, a nonzero deflection is present when *F* = 0 and *L*_*e*_ is non-zero and may become substantial as *L*_*e*_ becomes large.

To offset unloaded deflection from the variable spring weight and the fixed weight of the treadmill, a pretensioned torsion spring assembly was coupled between the end of the leaf spring and the ground. This assembly is designed to provide a pseudo-constant preload force as the device deflects under load. This was accomplished by preloading the springs such that a relatively small range of their full stroke is driven by the range of motion of the device. The vertical force applied to the leaf spring from each set of jaws of this preload device can be characterized as

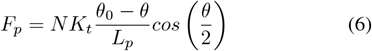

where *K*_*t*_ is the torsional spring stiffness, *N* is the number of torsion springs in parallel, *θ*_0_ is the rest angle of the torsion springs, and *L*_*p*_ and *θ* are the link length and angle between the links (Fig. 2B).

The angle between the links can be defined with mechanism geometry parameters

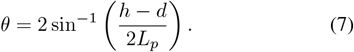

where *h* is the vertical distance between the mechanism connections the treadmill and to the ground at *d* = 0.

Substituting Eqn. 7 into Eqn. 6 and setting *θ*_0_ = 2*π* to indicate a maximum torsional deflection of one full revolution for the selected springs, the full equation for the preload force is

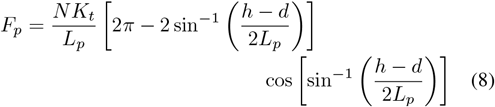

which can be simplified to

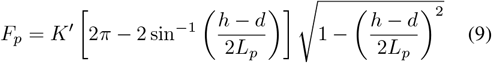

where *K*^*′*^ represents 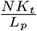 as the combined linear stiffness term, the second term represents the spring deflection in radians, and the third term represents the alignment of the resultant force with the movement axis of the treadmill as a unitless scalar bounded between 0 and 1.

For small (*h* − *d*)*/*2*L*_*p*_, the torsion spring deflection scales approximately linearly with the vertical deflection of the treadmill. The resultant force alignment scalar is more sensitive to this ratio, but nonlinear contributions from force alignment can be outweighed by large contributions of starting preload force.

In practice, the deflection compensation mechanism was implemented with two linkage sets nested in parallel to achieve sufficient upward force within the geometric constraints of our design. Eqn. 9 expresses *F*_*p*_ as a function of the treadmill deflection, subject to static geometric parameters of each linkage. The resultant calculation for *F*_*p*_ with multiple linkages is therefore the sum of Eqn. 9 calculated for each linkage for a given *d*. The implemented geometry can be competently described as a preloaded linear spring system, as confirmed by a linear approximation of preload force

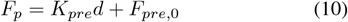

calculated via least squares regression. The linear approximation remains within 1% of the model defined by Eqn. 9 for the parameters of our mechanism.

By adding Eqn. 10 to Eqn. 5, the required external force for a given deflection can be expressed as

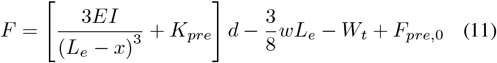

and the expected unloaded deflection is

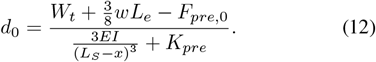

The vertical stiffness of the full assembly can be thus represented by

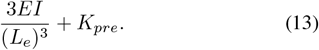

Actual values for the design parameters defined above are reported in Table I for the adjustable stiffness mechanism, and in Table II for the preload mechanism. The vertical stiffness is modeled to range between 5.2–3100 kN/m for these parameters. Unloaded deflection is modeled to range from 0– 39 mm for this range, with deflection exceeding 10 mm below a stiffness setting of 28 kN/m. Without the compensation mechanism, maximum unloaded deflection is modeled to reach 81 mm and exceed 10 mm below 56 kN/m.

**TABLE I.**
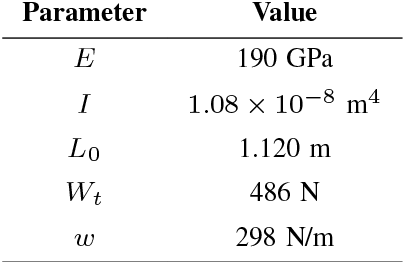
Adjustable stiffness design parameters

**TABLE II.** Preload mechanism design parameters

| Parameter | Linkage 1 | Linkage 2 |
| --- | --- | --- |
| $N$ | 12 | 9 |
| $L_p$ | 0.2 m | 0.3 m |
| $h$ | 0.295 m | |
| $K_t$ | 0.77 Nm/rad | |

The remainder of this section details the practical implementation of the design modeled above into a functional prototype.

### D. Design Implementation

#### 1) Mechanical Construction

The cantilever beam consists of three 4140 alloy steel plates suspended parallel to the ground by an aluminum base (Fig. 1). Two pairs of aluminum rollers mounted to a cart function as a moving cantilever base by constraining the displacement and curvature of the spring behind the lead roller pair. The cart rides on a linear track with ball bearings and is driven by a servomotor-actuated rack and pinion. The rack is 1 m long, allowing the effective length of the spring to vary from 0.12 m to 1.12 m. The minimum effective spring length is determined by the clearance required between the cantilever rollers and the attachments at the spring tip. The reaction forces from the spring are transmitted to the treadmill through a rigid link coupled to the tip of the spring and the base of the treadmill via steel shafts and needle bearings. These couplings function as revolute joints which allow the treadmill to deflect vertically and the spring tip to deflect through an arc without locking the mechanism.

The preload mechanism beneath the treadmill is constructed from a linkage machined from aluminum 6061 plates. The linkage is coupled to the spring tip attachment shaft with needle bearings and to the ground-fixed structural frame via an identical shaft and set of needle bearings. A total of 21 torsion springs with 0.77 N-m/rad stiffness each are mounted in parallel within this mechanism at an average preload of 78% of their maximum allowable deflection (Table II), resulting in a vertical force of 266 N under no additional load, assuming no friction losses. The preload mechanism is equipped with pins and hardstops to limit the maximum vertical deflection of the treadmill to be flush with the height of the adjacent rigidly mounted treadmill.

The treadmill, which has a mass of 49.6 kg (55% of which is counteracted by the preload mechanism), is constructed from aluminum U-channels and sheets. The top surface is lined with low-friction phenolic wear plates. The belt tension can be adjusted via a lead screw mechanism which positions the front roller. The treadmill is constrained to only move in the vertical direction via a set of linear bearings and support-rail shafts. The linear motion system is designed with up to 10 cm of allowable vertical travel, the maximum downward limit of which may be reduced in 1.3 cm increments by a set of telescoping pylons which function as hardstops. These pylons also include hardstop upward travel limits, redundant with the mechanical limits included in the preload mechanism.

To avoid restricting a user’s natural step length or other gait kinematics, the treadmill was designed with an effective walking surface length of 1.68 m. This is significantly longer than other elastically-suspended treadmills; therefore, additional measures must be taken to prevent the inertia of the treadmill from reducing the natural frequency of the system below that required for walking. We used two approaches to limit the treadmill inertia. First, the drive motor for the treadmill belt is located offboard. This is critical because the treadmill drive motor has a mass of 43 kg, which is nearly equal to the mass of treadmill itself. This motor drives the belt as it deflects vertically by coupling to the rear roller through universal joints. Second, unloaded deflection is counteracted by the spring powered preload mechanism described previously rather than by an inertial counterweight.

The treadmill is surrounded by a wooden platform to provide a rigid surface to safely step onto and mitigate the perception of balancing on a moving surface high above the ground. This platform is located 13 cm below the treadmill top surface. The treadmill may therefore sink nearly to the surrounding ground level at maximum deflection. Furthermore, the platform encases the adjustable stiffness mechanism for added safety.

#### 2) Actuator Configuration and Control

The position of the cantilever which defines the surface stiffness is controlled by a servomotor powering a rack and pinion actuator. While similar systems have been developed using a lead screw actuator [20], [21], [28], the screw length required for the target stiffness range would likely cause detrimental transverse vibration (i.e., “screw whip”) for the high rotation speed required to complete a maximal stiffness change within one step, except for very large screw diameters. The length of a rack and pinion system does not affect its performance if the rack is stationary, with the tradeoff that the inertial load the motor drives includes its own mass.

The servomotor was selected by calculating the maximum horizontal force the motor must exert and the time in which the largest stiffness change must be achieved (*<* 500 ms). The maximum force the motor must overcome to move the cantilever occurs at the stiffness setting where the vertical displacement caused by the applied downward force on the treadmill is equal to the mechanical deflection limits. Under this condition, the spring maximally deflects while transferring all of the load to the movable cantilever base. The horizontal force can be estimated using a formula that has been previously derived for cantilever beam deflection [30], [32]:

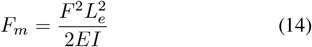

Assuming a maximum vertical load of 1.5 body weights of the heaviest allowable user (150 kg) and including a safety factor of 1.5, this force is estimated at 882 N. Therefore, we selected a 1kW servomotor (SV2L-210B, AutomationDirect, Cumming, GA) and 1.5 kW drive (SV2A-2150, AutomationDirect, Cumming, GA) with an attached 5:1 ratio gearbox and 24-bit incremental encoder to drive the system. This actuation system is rated to move the cantilever position at a maximum speed of 2.35 m/s with a force of 1100 N intermittently, or 430 N continuously. Assuming that the motor could reach maximum speed within 100ms, we estimated that these performance specifications would allow a change from maximum to minimum stiffness (or vice versa) in under 500 ms with an applied load on the treadmill up to our designed maximum. To further minimize demands on the motor, an integrated brake engages after the desired position is reached.

Stiffness control is performed on the servo drive via position and velocity trajectory tracking with a PI control architecture and 8 kHz sample frequency. Because stiffness can be mapped to effective spring length and does not require active control to maintain once set, static position targets were stored in the drive corresponding with the values characterized in the following section. The stiffness setting may be selected and triggered by controlling the state of corresponding digital inputs. Position targets and velocity profiles may be reprogrammed if modifications are made to the design or if more gradual change in stiffness is desired. Mechanical limit switches and hard stops mounted to a linear guide shaft prevent the cantilever cart from exceeding its designed range of motion and provide a reference position for homing the motor encoder should the zero position be lost. The system is further equipped with an emergency stop located at the operator workstation. System re-homing and stiffness control commands may be triggered by an operator using a custom pendant control panel.

A 3.7 kW AC motor (Leeson Electric Corporation, Grafton, WI) drives the treadmill belt via a 2.22:1 timing belt pulley, allowing a maintained maximum belt speed of 3.25 m/s with up to 1100 N of propulsive or braking force applied to the belt by the user. The belt speed is controlled with a 16-bit microprocessor based AC motor drive (Leeson Electric Corporation, Grafton, WI). An identical motor and drive power the belt for the adjacent rigid treadmill.

## IV. EVALUATION

To characterize the dynamics of the system and evaluate its ability to function as an experimental device, we performed a series of tests. The evaluation tests were designed to (1) quantify the actual stiffness behavior and range of the treadmill, (2) the natural oscillation frequency and damping ratio of the treadmill at different stiffnesses, and (3) the speed of the maximal stiffness change under load.

### A. Stiffness Characterization

To quantify the relationship between surface stiffness and effective spring length, we measured vertical deflection under static loads at 6 cantilever positions across the available range of spring lengths. We sequentially stacked and removed exercise weights on the treadmill belt up to 99 kg (971 N). We recorded the vertical position of the treadmill for each change in weight so as to observe the linearity of the stiffness relationship and capture any hysteresis effects. We measured the vertical position of the treadmill surface by applying six reflective markers at the four corners and two long-edge midpoints of the walking surface and recorded their positions with the motion capture system mentioned in Section III recording at 100 Hz. Each deflection measurement represents the vertical height of the markers averaged across three seconds (300 frames of data), and averaged again across all six markers. We discarded measurements for which the treadmill stopped against the mechanical limits, which occurred only for the minimum stiffness condition.

For each spring length, we fit a linear regression model to applied force vs. displacement, as illustrated in Fig. 3A. Using the slope of each linear fit as the approximate linear stiffness, we calculated the relationship between the stiffness and effective spring length (Fig. 3B). We recorded a maximum stiffness of 280 kN/m and a minimum stiffness of 8.6 kN/m, with all linear approximations achieving *R*^2^ values exceeding 0.97 except for the minimum stiffness condition, in which the *R*^2^ was 0.75, due to hysteresis.

**Fig. 3.**
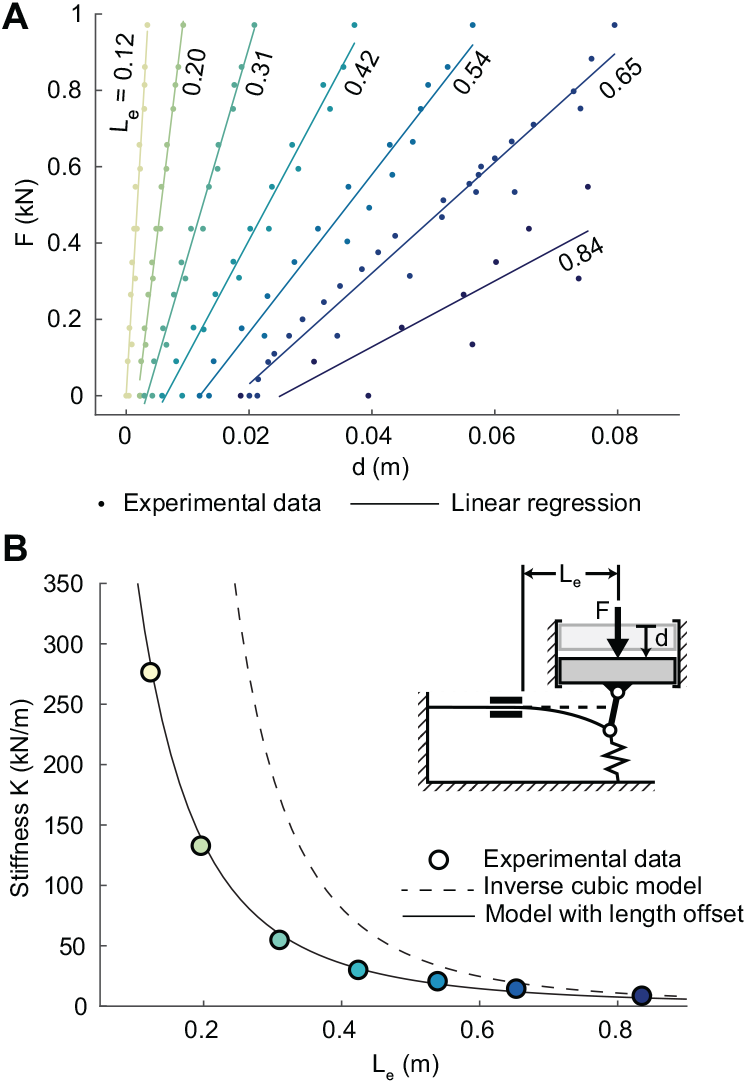
**A:** Linear stiffness of the adjustable stiffness mechanism was characterized at several position settings by recording treadmill displacements under loading with discretely applied weights. We estimated the linear stiffness coefficient by performing least squares linear regression for each setting and recording the slope of the resulting line. **B:** Surface stiffness vs effective spring length using the corresponding linear fits in **A**. The model developed in Section III is depicted with a dashed line. The same model with an offset of 0.16 m added to ***L***_***e***_ is depicted with a solid line, and fits the experimental measurements with an ***R***^**2**^ of 0.999, indicating that the cantilever constraint is not ideal.

Deflection of the walking surface without added load remained below 20 mm for most conditions, but it reached 21 mm before loading and 39 mm after loading at minimum stiffness. In deriving an expression for treadmill stiffness with respect to cantilever position, the original model overestimated the stiffness at short spring lengths. However, after applying an offset of 16 cm to the effective spring length and maintaining all other parameters, the measured stiffness closely followed the relationship predicted by the model (*R*^2^ = 0.999) (Fig. 3B). Note that we proceed to use the 15 kN/m setting as the minimum stiffness setting in all further tests due to the high degree of hysteresis, the magnitude of unloaded deflection, and the likelihood of the treadmill displacement being constrained under the test loads by the mechanical stops at 8.6 kN/m. We also approximate the maximum stiffness condition as ∼300 kN/m due to measurement precision, explained further in Section V.

### B. System Identification

To evaluate the speed of the passive device dynamics relative to the frequency of human walking, we applied a step change in vertical load to the treadmill at 300, 30, and 15 kN/m and measured the transient response of the vertical displacement of the walking surface using the same marker and camera setup as from the stiffness characterization evaluation. We approximated a step change in downward force by having a volunteer (mass = 68.9kg, or 676 N) step onto the treadmill from a waiting position with one foot held over the walking surface. To account for variability in the load application, we repeated this procedure four times for each stiffness condition and averaged the treadmill response, synchronized to when the belt position deflected by 1 mm.

To quantify the system dynamics, we approximated the response as a second-order spring-mass-damper system. The second-order model was fit to the data via iterative error minimization using the Matlab system identification toolbox (Mathworks, Natick, MA). Model fit assessed from 0 to 100% as normalized root mean squared error (NRMSE) was 94.7%, 93.6%, and 94.4% for the 300 kN/m, 30 kN/m, and 15 kN/m conditions, respectively. The actual response and modeled second-order response are illustrated in Fig. 4A. From high to low stiffness, the natural frequency was 3.2, 2.4, and 2.1 Hz, all of which exceed the average stride frequency of walking (approx. 0.9–1.1 Hz, [33]). The system is underdamped, with damping ratios of 0.48, 0.30, and 0.27 from high to low stiffness. Extrapolating the frequency response of these second-order models (Fig. 4B), phase lag is less than 20° at typical walking cadence. Response magnitude falls below -3 dB at 4.1, 3.5, and 3.1 Hz from high to low stiffness.

**Fig. 4.**
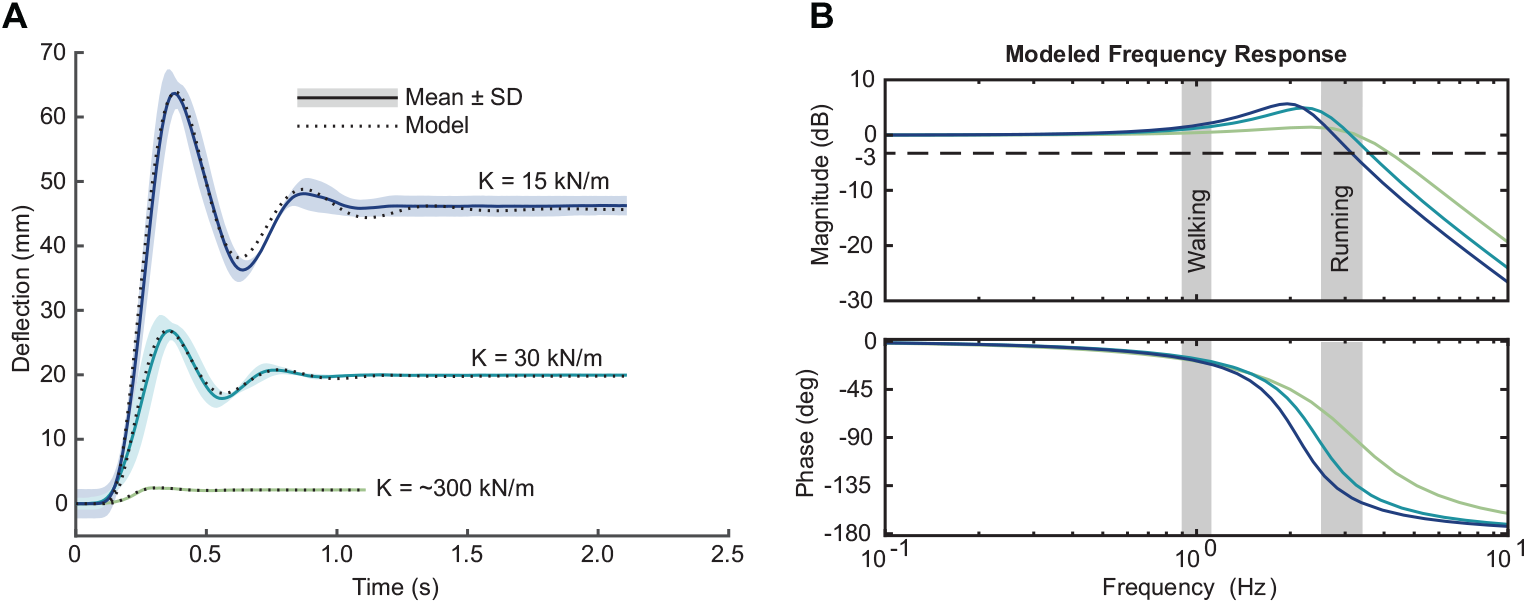
**A:** Transient response to a step input of 676 N and the modeled response of a second-order system. The second-order model was fit to the data via iterative error minimization using the Matlab system identification toolbox. Model fit assessed from 0 to 100% as normalized root mean squared error (NRMSE) is 94.7%, 93.6%, and 94.4% for the 300 kN/m, 30 kN/m, and 15 kN/m conditions, respectively. **B:** Bode chart of the frequency response for the modeled second-order systems. Shaded regions illustrate the input frequency range expected during walking and running. Note that the walking frequency range is based on the stride frequency, as the treadmill is intended interact with only one leg during walking gait. The running frequency range is based on the step frequency, as footfalls occur in-line during natural running gait, requiring both feet to land on the same belt.

### C. Stiffness Control Performance

To evaluate the active control of the adjustable stiffness, we applied position control trajectories between the 15 kN/m to 300 kN/m settings with and without a load disturbance applied to the treadmill in the form of a person standing on the belt (80.7 kg, or 792 N). Rather than applying step changes to the reference position, we applied trapezoidal trajectories of 450 ms duration with smooth acceleration and deceleration for the initial and final 40 ms to minimize peak current demands on the motor. We recorded cantilever position and velocity from the embedded motor encoder and motor current as reported from the motor driver at a frequency of 8 kHz. Each position command was repeated 6 times.

We calculated the mean position, velocity, and motor current for increasing and decreasing stiffness change with and without added weight to the treadmill (Fig. 5). Note that the 90% rise time is unaffected by the presence of a load disturbance and remains at ∼340 ms for all conditions.

**Fig. 5.**
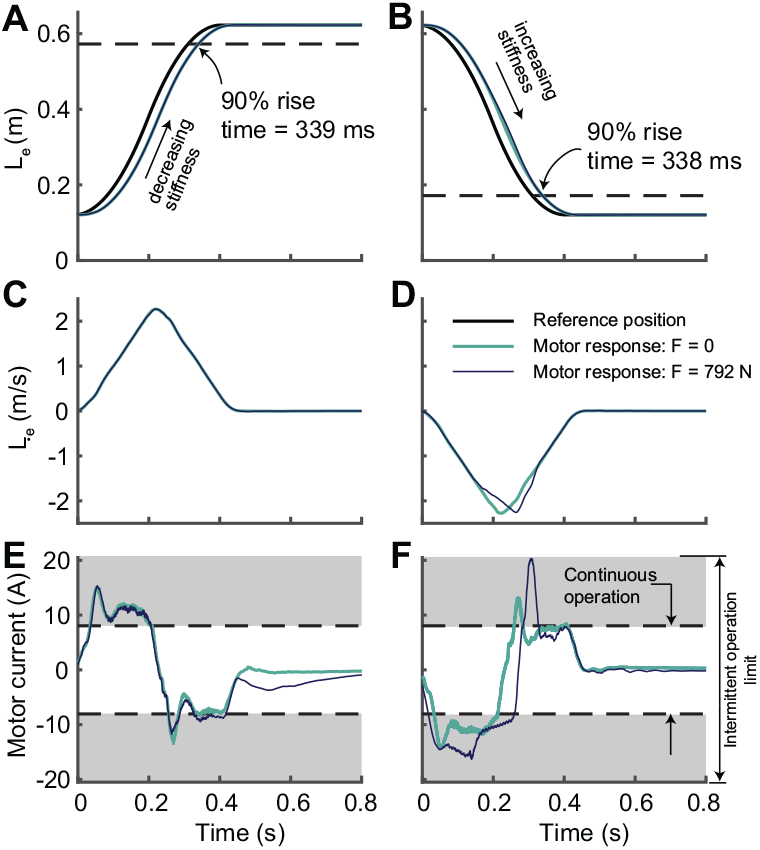
Motor control performance increasing and decreasing stiffness between 300 kN/m and 15 kN/m with and without an applied load to the treadmill. **A:** Effective spring length for decreasing stiffness and **B:** increasing stiffness. **C:** Rate of change of effective spring length for decreasing stiffness and **D:** increasing stiffness. **E:** Electric current drawn by the motor to track the trajectories in **A** and **C. F:** Electric current drawn by the motor to track the trajectories in **B** and **D**.

### D. Operation During Walking

Finally, to qualitatively demonstrate the behavior of the system in a practical human experiment scenario, we recorded the vertical treadmill position during a stiffness change from 300 to 30 kN/m while a person (74.4 kg) walked on the belts at 1.25 m/s (Fig. 6).

**Fig. 6.**
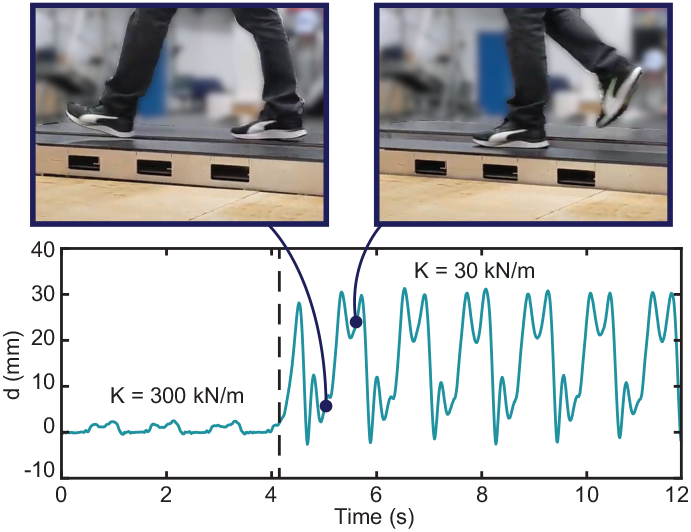
Vertical deflection of the treadmill during 1.25 m/s walking. Stiffness changes from 300 kN/m to 30 kN/m within one step at t = 4.1s.

## V. DISCUSSION

We developed a treadmill and suspension system capable of adjusting the surface stiffness within a range from 15 to 300 kN/m of well characterized linear stiffness during a single step, swing or stance phase. The AdjuSST meets these requirements while maintaining an effective walking surface length of 1.68 m, more than double the step length of a person of average height, and deflecting vertically downward up to 10 cm. The natural frequency of the AdjuSST exceeds the input frequency of walking at all available stiffnesses. Therefore, the AdjuSST successfully achieved our proposed contributions.

### 1) Stiffness characterization

Measuring the static displacement of the treadmill in response to fixed loads demonstrated that the effective vertical stiffness is linear across the available range. We observed increasing hysteresis as stiffness decreased. This is likely due to static friction in the mechanical suspension resisting rebound of the treadmill surface for incremental reductions in vertical load. Additionally, while the stiffness behavior is well characterized and offers a useful range, the cantilever constraints at both the moving cart and the fixed base are not ideal and result in a smaller upper bound on stiffness than we anticipated. This is most likely due to the relatively short length of the constraints and the presence of small clearances between cantilever rollers and the spring, which combine to allow the spring to bend upward behind the constraint.

We report the maximum stiffness as a rounded approximation because the calculation of high stiffness becomes sensitive to the resolution of the displacement measurement. The motion capture camera system was calibrated to within 0.5 mm of error, and the largest deflection measured at maximum stiffness was 3.4mm, meaning that the maximum stiffness could plausibly be a value between 250 and 340 kN/m.

### 2) Passive suspension dynamics

Identification of the system dynamics revealed that the treadmill suspension has sufficient frequency bandwidth and resilience (i.e., not dominated by energy dissipation) to present a spring-like response to walking gait without encountering substantial phase lag or failure to return to the undeflected position before the next heel strike. Assuming running cadence is between 70 and 110 strides per minute [34], [35] and that both feet interact with the same belt, input frequency during running would vary between 2.3-3.7 Hz (Fig. 4B). Thus, the system response is likely to be distorted for low stiffness at high running cadences, but it may near resonance in the higher stiffness range at slow running cadences. This resonant frequency band may be useful in running contexts, where the objective of changing stiffness is more likely to align with improving energy economy by taking advantage of the foot-ground interaction dynamics [24], [36], [37] than with the asymmetric stiffness perturbation paradigm the treadmill was originally designed to enable. Finally, we found that the treadmill response is underdamped, which provides a useful baseline for potentially studying the effects of added damping in the future.

### 3) System actuation

The actuation performance tests demonstrated that the AdjuSST can change between stiffness extremes within one step, irrespective of the walker’s weight on the treadmill. For the tracking profile tested, motor current remained near the rated continuous operation limit during the stiffness change. The maximum rated intermittent current is briefly reached only when decelerating at the end of an increasing stiffness change with the disturbance load applied. Though the rated continuous current was exceeded, the system will not be continuously actuated, and will at most be actuated twice within one second with long idle periods in between when performing single step perturbations. Note that the system is also capable of following more gradually changing trajectories at reduced demand on the motor.

### 4) Limitations

Despite the successful contributions of this design, our approach has some limitations. Despite our countermeasures, some deflection at zero added load is present for lower stiffnesses because the preload mechanism does not compensate the full weight of the treadmill. While we have limited this deflection to under 1 cm across most of the stiffness range, it becomes noticeable at the extreme low end. This deflection remains for two main reasons: (1) The steel spring is substantially heavy that fully eliminating zero-load deflection will create a “dead zone” at higher stiffnesses, where a noticeable preload must be overcome before the belt will begin to deflect downward, and (2) the number of springs or jaws required to eliminate zero-load deflection is limited by the geometric constraints of our design. The design presented is a compromise between surface uniformity, treadmill size, and the ability to present an instantly reactive surface. The preload mechanism design is flexible enough to allow this operating point to be shifted toward increased preload if desired by the addition of higher stiffness torsion springs. This design also cannot reach the high range of stiffness achieved by the VST because the cantilever constraint cannot bypass the spring in either direction, as can be done by locating the fulcrum point of a rigid lever directly at one of the load insertion points. We chose to accept this limitation because our target range was achievable with this design, and it comes with other benefits, such as requiring less vertical height for the suspension compared to other methods and requiring low startup torque from the motor under heavy loads.

### 5) Comparison with other adjustable surface stiffness treadmills

As previously mentioned, two other adjustable surface stiffness treadmill designs have been presented [20], [21]. Evaluation metrics that can be directly compared between designs are included in Table III. Perhaps the clearest distinction between the AdjuSST and other designs is the physical dimensions: the effective walking surface is over twice as long as the VST and sits lower to the ground than either of the other designs. As stated in the above paragraph, the stiffness range of the AdjuSST is narrower than the VST, but AdjuSST achieves significantly higher stiffness than the TwaS. Like the TwaS, the AdjuSST deflects linearly rather than angularly about a fixed axis like the VST. The belts of the AdjuSST are driven by motors rated for 5*×* more mechanical power than the VST, allowing it to run at nearly twice the maximum belt speed while also overcoming higher braking/propulsion forces to maintain that speed. The drive motors of the TwaS are unspecified, though are reported to run the belts at up to 3.6 m/s, which is slightly faster than the AdjuSST at its current drive pulley ratio. All three designs are reported to be able to complete a maximum stiffness change in under 500 ms, though the VST is the fastest, with rise times under 250 ms. Unfortunately, a direct comparison of the system dynamics of the spring suspension is not possible, as this evaluation was not performed with the other designs. The open-loop response of the VST stiffness change was characterized at a similar bandwidth to the AdjuSST suspension, but these measures are not perfectly analogous.

**TABLE III.** Comparison of Adjustable Stiffness Treadmills

| Parameter | AdjuSST | VST [20], [23] | TwAS [21] |
| --- | --- | --- | --- |
| Platform height (m) | 0.67 (0.13)* | 0.70 | 1.04 |
| Walkable length (m) | 1.68 | 0.8 | - |
| Stiffness range (kN/m) | 15–300 | 0.06–1000 | 0.05–40 |
| Linear motion (Y/N) | Y | N | Y |
| Suspension $\omega_n$ (Hz) | 2.1–3.2 | - | - |
| Suspension $\zeta$ | 0.27–0.48 | - | - |
| Max. belt speed (m/s) | 3.25 | 1.85 | 3.6 |
| Belt motor power (kW) | 3.7 | 0.8 | - |
| Stiffness rise time (ms) | 339 | < 250 | ~ 500 |

\* Indicates distance to raised deck
Omitted values are not reported in cited manuscripts

## VI. CONCLUSIONS

This paper presented and evaluated a novel adjustable surface stiffness treadmill design. Overall, the evaluation of the AdjuSST has demonstrated that it is a capable platform for conducting experiments involving asymmetric perturbations of surface stiffness. In context with alternative designs, the AdjuSST contributes meaningful improvements in effective walking surface length while maintaining a useful stiffness range and responsive spring suspension. Future work with this system will focus on experiments which perturb human gait with prolonged asymmetric stiffness changes to investigate adaptation effects and aftereffects relevant to neuromotor rehabilitation.

## Acknowledgment

We thank the many undergraduate and graduate students who contributed their time and effort toward building this system: Aidan Downey, Samuel Naftal, Simon Houghton, Alexander Klinkhamer, Kyle O’Connell, Emily Brann, Timothy Chu, Onur Bulut, Deanna Conti, Maria Hernandez, Robert Bennett, and Molly Fabrizio. We thank Jenna Chiasson, Leah Metsker, Elena Schell, and Calder Robbins for their assistance during validation experiments.

